## Supplementary for "Targeted Computational Design of an Interleukin-7 Superkine with Enhanced Folding Efficiency and Immunotherapeutic Efficacy"

**Supplementary Data**

**(A)**

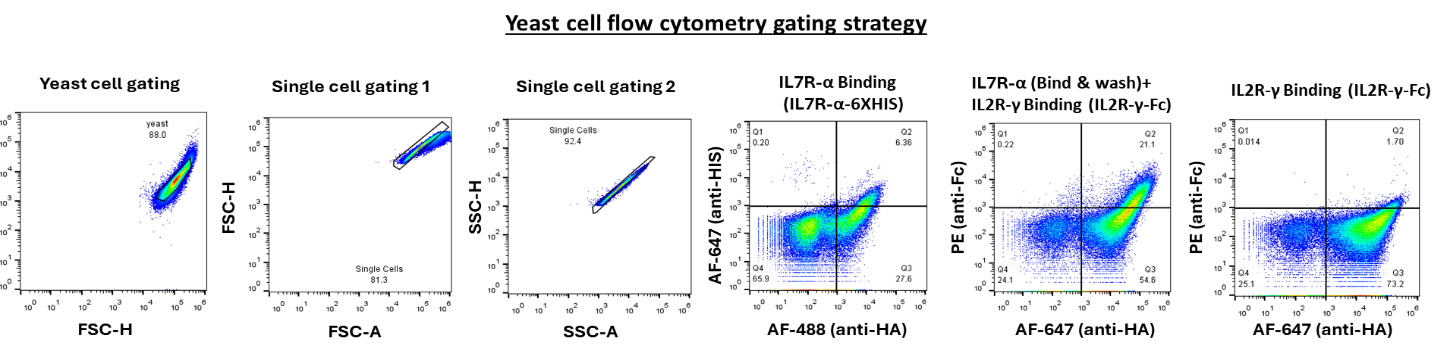

**(B)**

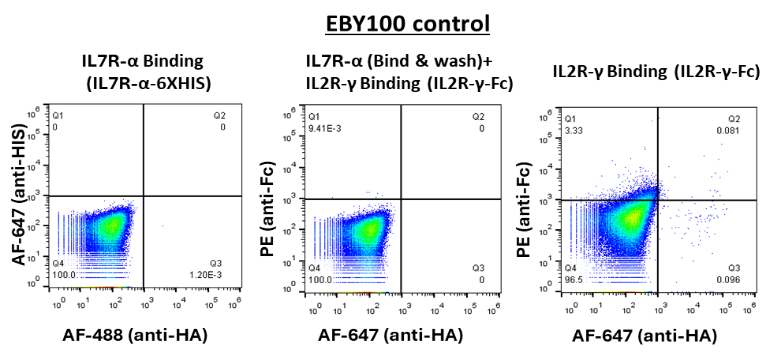

**(C)**

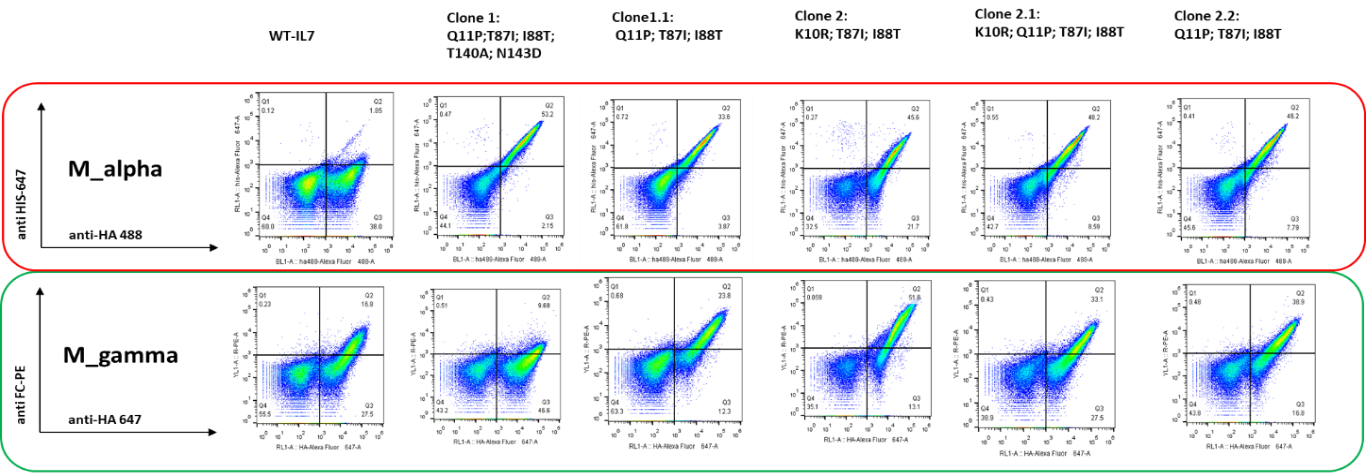

**(D)**

**Figure 3-figure supplement 1. (A) Flow cytometry gating strategy of yeast-display recombinant IL7 or Neo7s. The displayed protein (IL7/Neo7s) carries a HA-tag while the recombinant IL7 receptors carries either a HIS (IL7R-α) or a FC-tag (IL2R-γ). The signal intensity of X-axis (confers by binding of anti-HA mab) correlates with the expression level of the displayed protein while the signal intensity of Y-axis (confers by binding of anti-HIS/anti-FC mab to the recombinant receptors bound to the displayed proteins) correlates with the binding affinity of the displayed proteins towards IL7 receptors. IL7 binds to IL7R-α at high affinity followed by the dimerization with the IL2R-γ. IL7 and Neo7s bind poorly to the IL2R-γ in the absence of IL7R-α. Therefore, pre-incubation of the displayed protein with the IL7R-a is necessary (align with its biological mechanism) for proper quantification of the binding of IL7/Neo7s to the IL2R-γ. (B) EBY100 negative control for flow cytometry analysis. (C) Binding data with mIL7R-α. (D) Binding data with mIL7R-α and mIL2R-γ.**

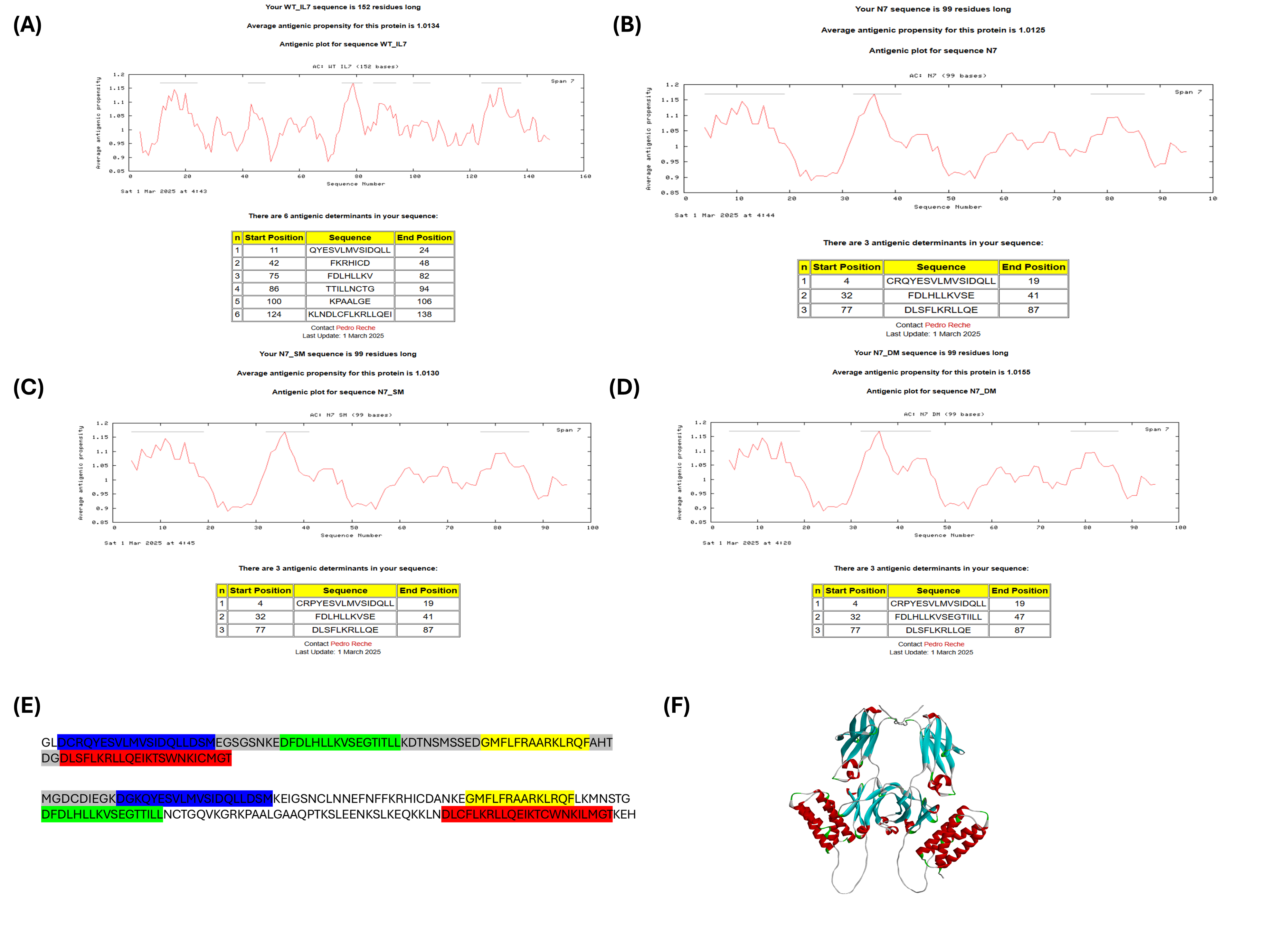

**Figure 5-figure supplement 1. Immunogenic epitopes prediction results from the sequence of (A) WT-IL7 (B) Neo-7 (C) Neo-7 with Q6P mutation (D) Neo-7 with Q6P and T45I mutation (E) Sequence of Neo-7 versus WT-IL-7, sequence of the identical helices were highlighted using the same color. The remodeled loops were colored as grey (F) Alphafold prediction of the structure of an Fc-Neo-7 fusion homodimer.**

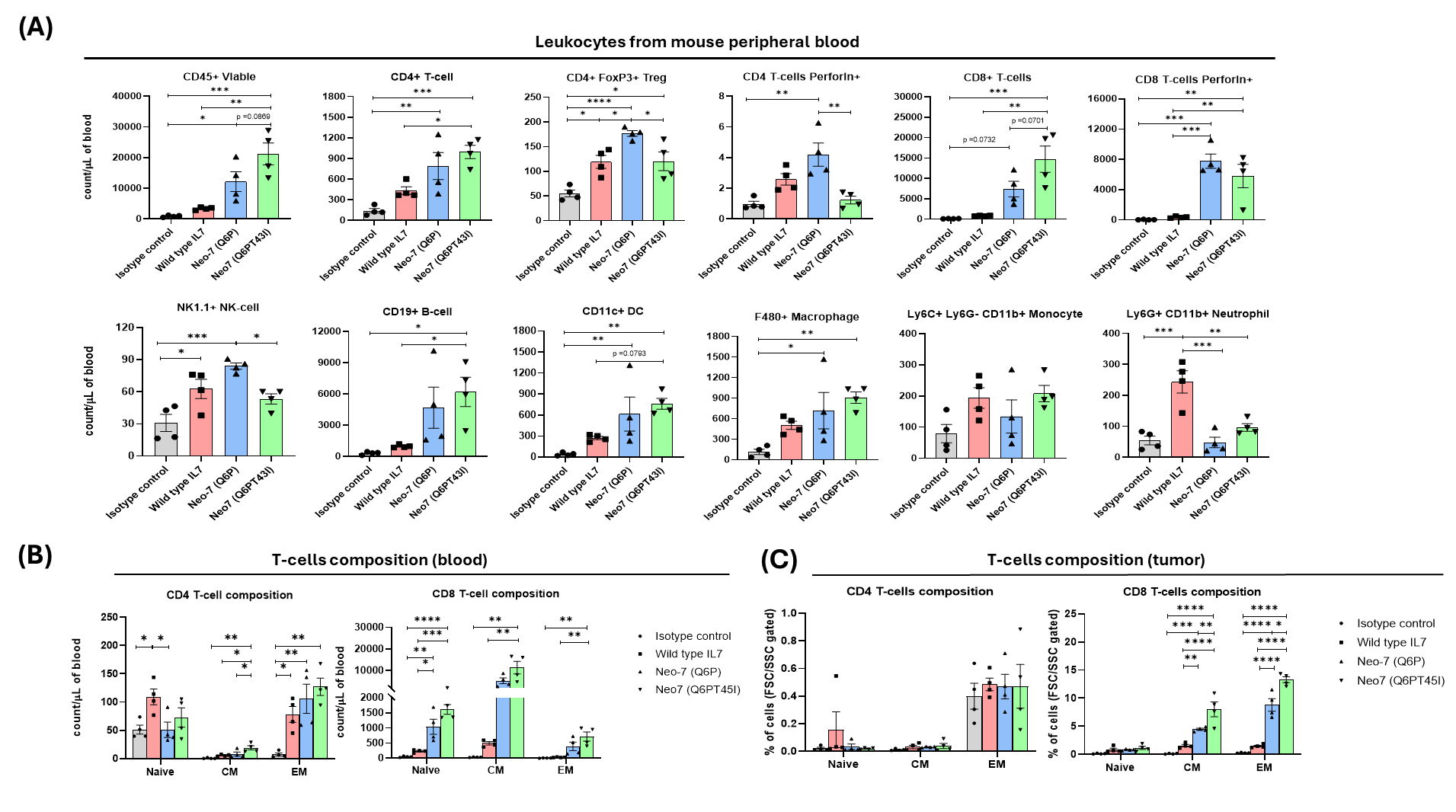

**Figure 7-figure supplement 1. (A) Counts of total leukocytes and different immune cell types within the PBMC of tumor-harboring mice at day 7 post-treatment. T-cell characterizations (B) in PBMC and (C) in tumor based on their expression of surface memory markers.**

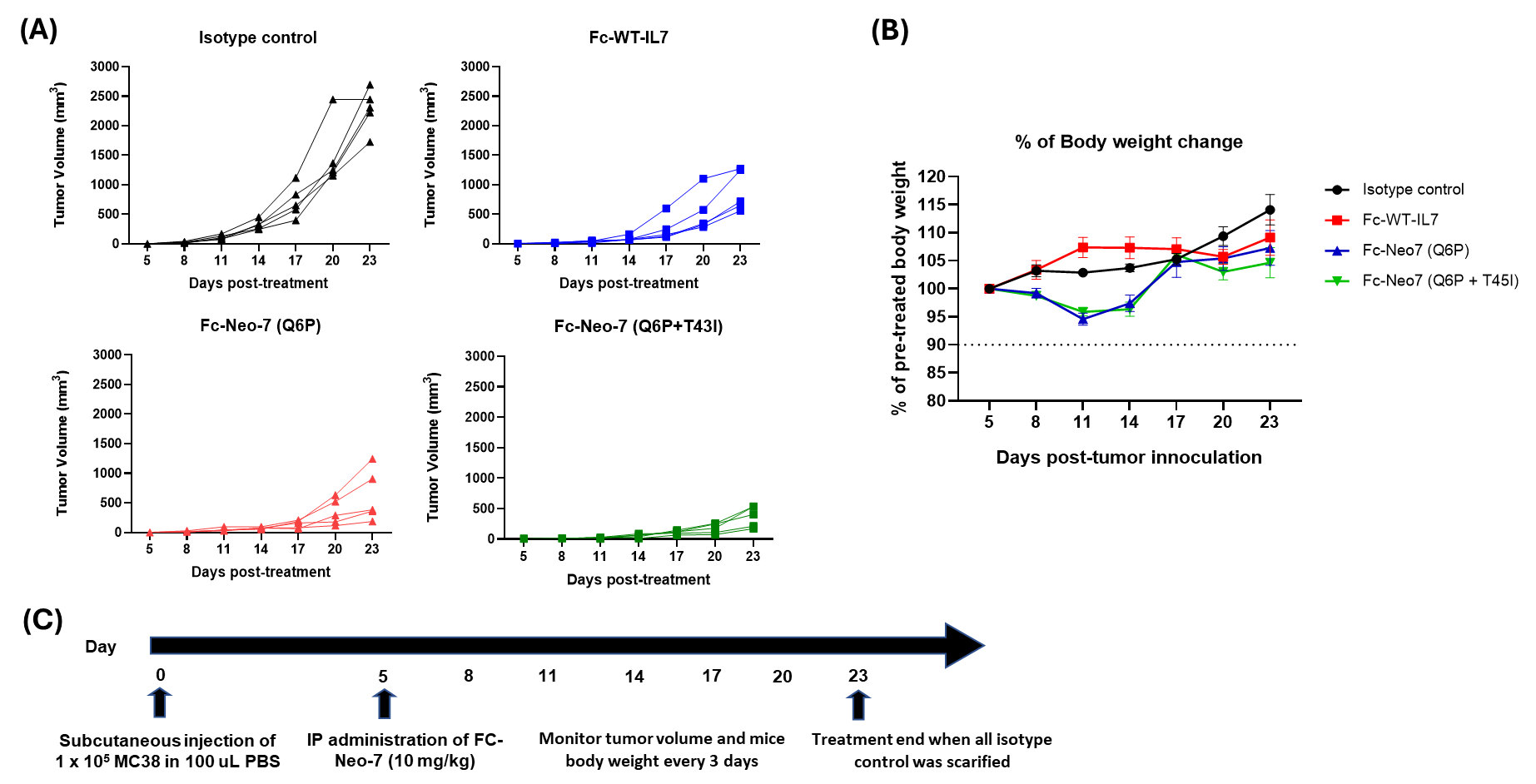

**Figure 7-figure supplement 2. (A) Individual tumor growth curve of mice in different treatment group. (B) Body weight of mice subjects in different group. (C) Dosing scheme of the *in vivo* anti-tumor assay.**

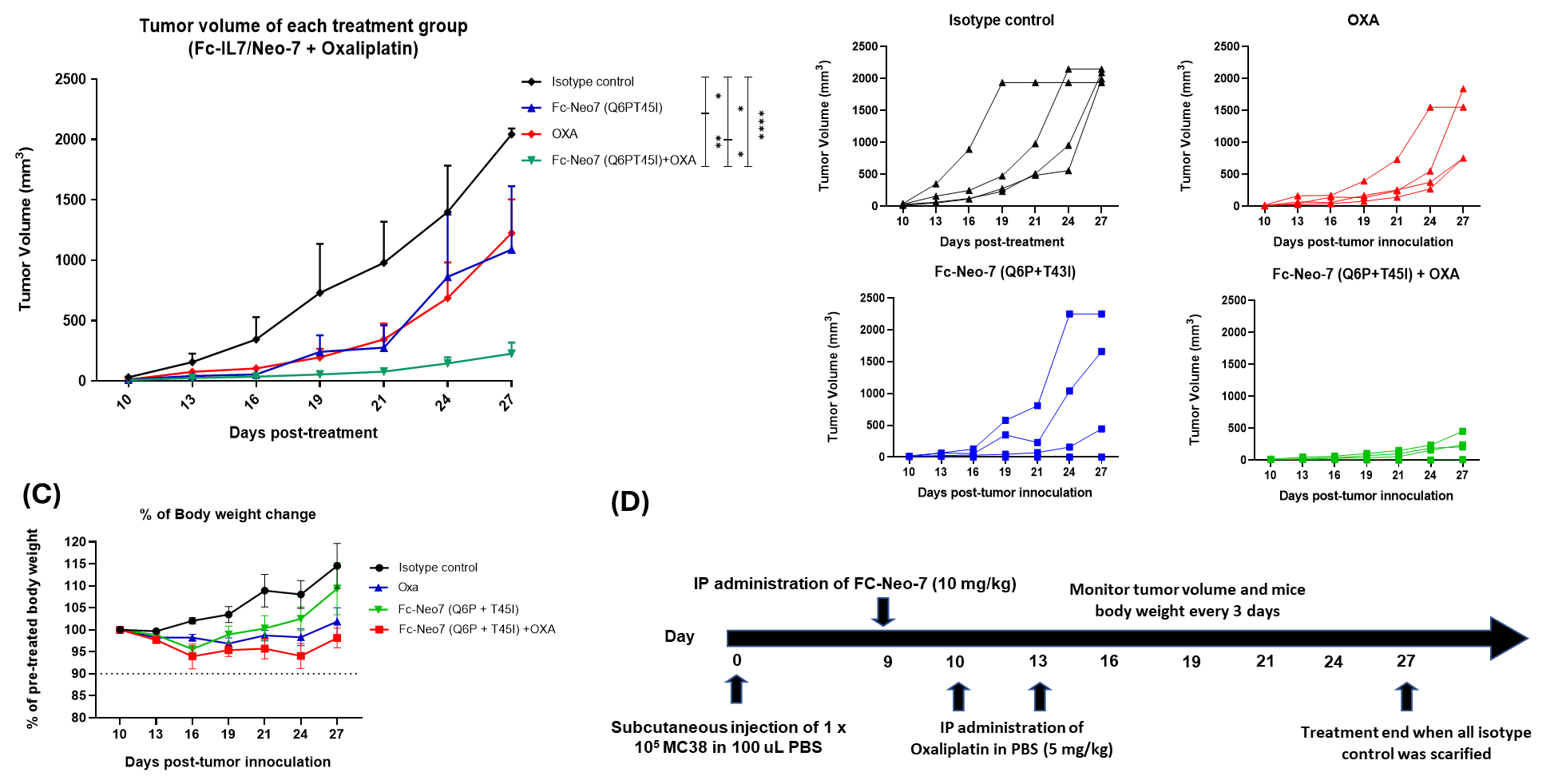

**Figure 7-figure supplement 3. (A) Tumor growth curve of MC38-tumor harboring mice received PBS, Neo-7, oxaliplatin or combination treatment as intervention. (B) Individual tumor growth curve of mice in different treatment group. (C) Body weight of mice subjects in different group. (D) Dosing scheme of the *in vivo* anti-tumor assay**

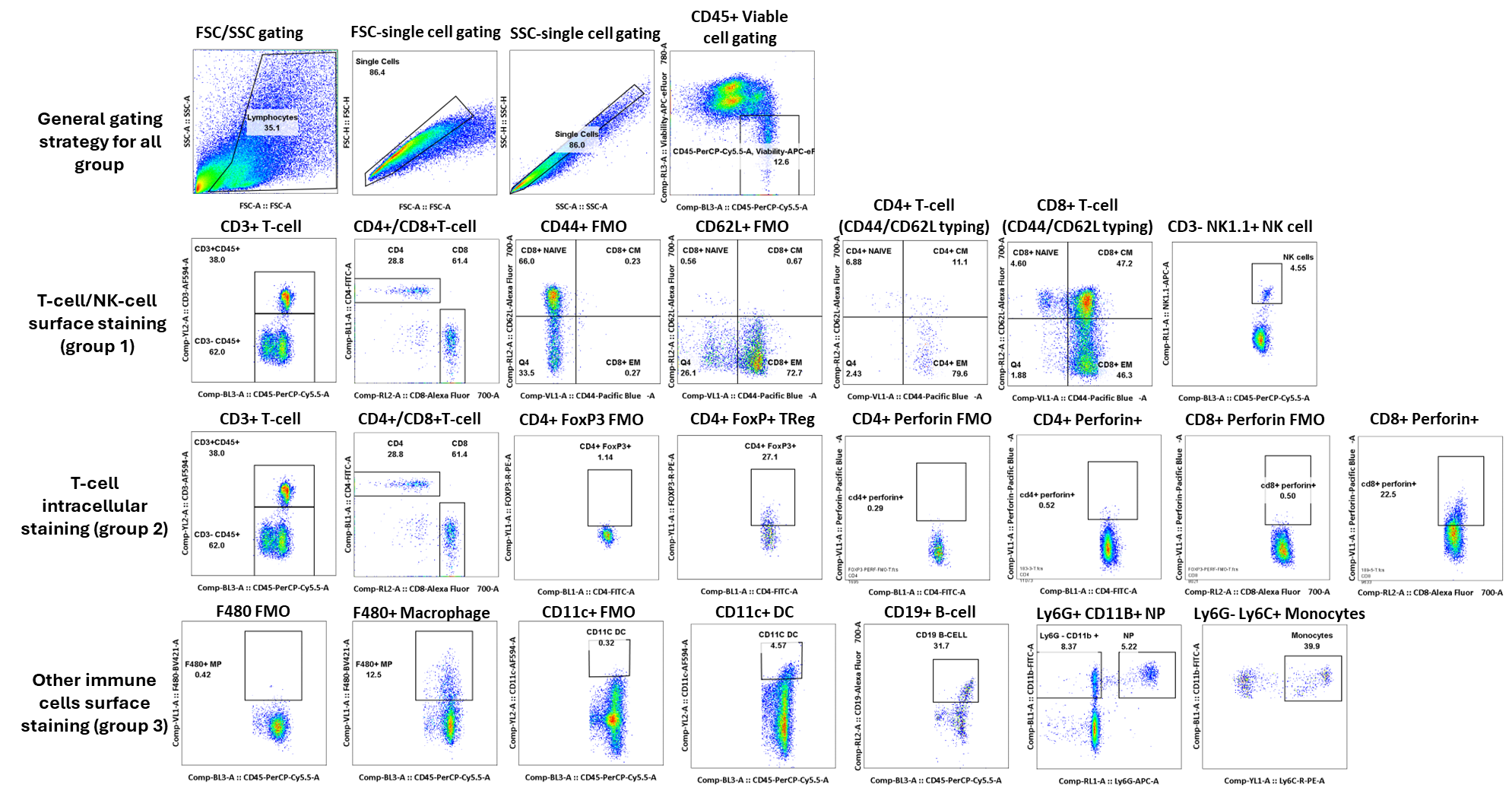

**Figure 7-figure supplement 4. Flow cytometry gating strategy of blood and tumor-extracted leukocytes in this study.** In general, all cells were gated by FSC and SSC to exclude debris, followed by single cell and the gating of CD45+ viable immune cells for further analysis. The leukocytes were divided into 3 staining groups. Group 1 includes the staining of NK-cell surface marker, T-cell surface and memory markers. Group 2 includes the staining of T-cell surface with intracellular staining of the FOXP3 and perforin markers. Group 3 consists of the surface staining of other immune cells such as macrophage, monocytes, neutrophils, B-cells and DC.

|  | **CD45+ CD3+** | **CD3+ CD4+** | **CD3+ CD8+** | **CD11C+ DC** | **CD19+ B-CELL** | **F480+ Macrophage** | **Ly6G- CD11b+ Monocyte** | **Ly6G+ CD11b+ NP** | **CD3+ CD4+ FoxP3+** | **CD3+ CD4+ PD1+** | **CD3+ CD4+ Perforin+** | **CD3+ CD8+ PD1+** | **CD3+ CD8+ Perforin+** |
| --- | --- | --- | --- | --- | --- | --- | --- | --- | --- | --- | --- | --- | --- |
| **FC-ctrl 1** | 485.87 | 31.32 | 62.65 | 24.39 | 147.19 | 285.48 | 103.95 | 40.35 | 44.77 | 21.84 | 1.91 | 17.50 | 10.03 |
| **FC-ctrl 2** | 537.32 | 49.46 | 46.25 | 36.68 | 34.52 | 324.37 | 184.01 | 122.93 | 8.42 | 13.11 | 1.00 | 3.82 | 1.04 |
| **FC-ctrl 3** | 426.80 | 23.54 | 39.48 | 65.04 | 33.04 | 327.09 | 246.05 | 73.82 | 2.49 | 3.85 | 0.49 | 7.02 | 1.37 |
| **FC-ctrl 4** | 447.41 | 41.98 | 35.31 | 77.28 | 61.48 | 320.74 | 238.52 | 91.85 | 17.78 | 22.04 | 1.48 | 23.89 | 15.37 |
| **WT-IL7-1** | 812.81 | 51.36 | 265.97 | 64.12 | 42.96 | 344.99 | 267.30 | 100.41 | 5.96 | 5.70 | 1.48 | 8.17 | 2.30 |
| **WT-IL7-2** | 880.28 | 41.26 | 269.61 | 71.58 | 55.02 | 490.11 | 254.88 | 100.35 | 16.00 | 10.42 | 2.95 | 142.53 | 79.26 |
| **WT-IL7-3** | 1347.39 | 74.23 | 498.74 | 131.17 | 42.52 | 410.81 | 271.35 | 40.00 | 23.51 | 14.32 | 1.35 | 14.32 | 2.16 |
| **WT-IL7-4** | 442.51 | 58.49 | 190.61 | 22.54 | 24.97 | 99.26 | 52.43 | 56.35 | 8.33 | 2.82 | 0.18 | 6.79 | 1.35 |
| **N7-Q6P-1** | 5693.51 | 132.07 | 3424.14 | 64.68 | 190.27 | 721.44 | 178.02 | 34.23 | 29.32 | 11.89 | 0.68 | 117.70 | 54.32 |
| **N7-Q6P-2** | 6294.12 | 152.35 | 3695.88 | 76.27 | 201.37 | 590.59 | 181.96 | 151.96 | 10.59 | 14.85 | 0.74 | 14.71 | 8.09 |
| **N7-Q6P-3** | 3435.45 | 126.18 | 2463.09 | 46.67 | 325.04 | 305.20 | 153.98 | 48.78 | 14.88 | 16.22 | 2.07 | 889.39 | 364.63 |
| **N7-Q6P-4** | 6236.99 | 182.80 | 4060.65 | 118.28 | 327.31 | 1071.40 | 303.23 | 54.62 | 46.13 | 45.81 | 1.29 | 1117.42 | 452.26 |
| **N7-Q6PT45I-1** | 9662.56 | 167.86 | 6798.29 | 101.54 | 295.04 | 724.79 | 223.59 | 33.85 | 36.92 | 36.41 | 5.13 | 1090.51 | 333.85 |
| **N7-Q6PT45I-2** | 6205.21 | 43.56 | 5590.60 | 39.24 | 98.16 | 299.68 | 105.65 | 31.24 | 6.95 | 8.76 | 0.57 | 85.43 | 122.29 |
| **N7-Q6PT45I-3** | 5110.91 | 60.00 | 2251.52 | 133.33 | 164.24 | 564.24 | 236.36 | 84.85 | 29.09 | 12.73 | 0.45 | 799.09 | 510.00 |
| **N7-Q6PT45I-4** | 5870.48 | 193.74 | 4328.44 | 58.23 | 110.75 | 397.28 | 117.82 | 40.27 | 48.98 | 25.71 | 1.84 | 499.80 | 202.24 |

**Figure 7-figure supplement 5. Absolute count of immune cells determined from TILs/mg of the tumor excised**
